# Characterization of the clade 4 non-toxigenic *C. difficile* isolate L-NTCD03 carrying the *cfr*(B) gene

**DOI:** 10.64898/2026.01.12.695941

**Authors:** Britt Nibbering, Sam Nooij, Céline Harmanus, Ingrid M.J.G. Sanders, Inez M. Miedema, Quinten R. Ducarmon, Rolf H.A.M. Vossen, Susan L. Kloet, Colleen K. Ardis, Robert A. Britton, Farnaz Yousefi, Jenna Bayne, Chandru Charavaryamath, Andy Law, Morgan L. Murphy, Brett Sponseller, Eric R. Burrough, Alejandro Ramirez, Shankumar Mooyottu, Tanja Opriessnig, Ed J. Kuijper, Meta Roestenberg, Wiep Klaas Smits

## Abstract

*Clostridioides difficile* infection (CDI) is a toxin-mediated gastro-intestinal disease. Yet, *C. difficile* is a phylogenetically diverse species that includes many non-toxigenic strains. In general, these are understudied, despite having significant potential impact for our understanding of the colonization process and as therapeutic modalities. Here, we present an in-depth characterization – including the complete genome sequence - of the non-toxigenic *C. difficile* strain L-NTCD03. This strain belongs to PCR ribotype 416, clade 4 and multilocus sequence type 39. It is resistant to multiple antimicrobials, but not those used for treatment of CDI. We validated the relevance of the *cfr*(B) gene from this strain in antimicrobial resistance to clindamycin, linezolid, retapamulin and streptogramin A. We found the L-NTCD03 strain to be non-toxic in various assays. Altogether, L-NTCD03 is a promising candidate for developing into a live biotherapeutic product.

## Introduction

The Gram-positive anaerobic spore-forming bacterium *Clostridioides difficile* is best known as a nosocomial pathogen and *C. difficile* infection (CDI) is regarded as an important cause of healthcare-associated infectious diarrhea (Leffler & Lamont, 2015, Smits *et al*., 2016). In this context, in particular strains belonging to the fluoroquinolone resistant PCR ribotype 027 (RT027) are associated with increased morbidity and mortality (Warny *et al*., 2005, He *et al*., 2013). Nevertheless, recent years have seen an increase in CDI cases that are not linked to classical risk factors such as antibiotic use or healthcare exposure, and are community-acquired (Smits *et al*., 2016, Viprey *et al*., 2022). Strains belonging to different types than RT027, such as those belonging PCR ribotype 078 (RT078), which also occur in many animal species, may be associated with community-acquired disease (Goorhuis *et al*., 2008, Viprey *et al*., 2022). Thus, *C. difficile* can be considered a quintessential One Health pathogen (Lim *et al*., 2020).

Considering its causative role in debilitating disease, much research has focused on the main virulence factors of *C. difficile:* toxin A (TcdA), toxin B (TcdB) and binary toxin (CDT) (Aktories *et al*., 2017, Kordus *et al*., 2022). Of note, whereas TcdA and/or TcdB is required for disease, CDT is mainly produced by a subset of toxigenic *C. difficile* strains, including those belonging to the epidemic types RT027 and RT078 (Smits *et al*., 2016).

*C. difficile* colonization does not always lead to disease. Importantly, a fraction of the population is asymptomatically colonized by toxigenic *C. difficile* (Crobach *et al*., 2018) and in particular in infants a high prevalence of *C. difficile* is noted without apparent disease (Ferretti *et al*., 2023, Mani *et al*., 2023). Additionally, *C. difficile* is not a universally pathogenic species; that is, non-toxigenic *C. difficile* (NTCD) strains exist that lack the genes encoding TcdA, TcdB and CDT, such as those belonging to PCR ribotype 010 (RT010) (Braun *et al*., 1996, Monot *et al*., 2015, Smits *et al*., 2016). Though these isolates have not been found to induce disease in humans and animals, it has been noted that they might act as reservoirs for antimicrobial resistance genes. For instance, resistance to metronidazole is common in RT010 strains (Boekhoud *et al*., 2020, Boekhoud *et al*., 2021) and over half the non-toxigenic isolates from Southeast Asia appeared to carry antimicrobial resistance determinants on mobile genetic elements (Imwattana *et al*., 2020).

*C. difficile* is a phylogenetically diverse species with isolates falling into at least 5 classical clades and 3 so called cryptic clades, that could be considered a separate genomospecies (Knight *et al*., 2021). Epidemic strains fall generally into clade 2 (RT027 and related strains) and clade 5 (RT078 and related strains). NTCD have been reported in multiple clades, such as clade 1 (which includes RT010)(Baktash *et al*., 2022), clade 4 (Imwattana *et al*., 2020), clade C-I and clade C-II (Ducarmon *et al*., 2023). Nevertheless, such strains are generally understudied compared to the toxigenic counterparts.

There is epidemiological and experimental evidence that NTCD carriage is associated with lower incidence of CDI, that NTCD colonization can prevent CDI disease and can reduce CDI recurrence in intestinal models, animals and humans (Natarajan *et al*., 2013, Gerding *et al*., 2018, Etifa *et al*., 2023). The strongest evidence for use of NTCD as a therapeutic intervention is for strain M3 (a RT010 isolate) in humans (Villano *et al*., 2012, Gerding *et al*., 2015, Sambol *et al*., 2023) and strain Z31 (RT009) in piglets (Oliveira Junior *et al*., 2019, Oliveira Junior *et al*., 2019).

In addition to potential use as a live biotherapeutic product, NTCD could provide a chassis for engineering of immune-epitopes and therefore serve as a potential vaccine (Hughes *et al*., 2022, Wang *et al*., 2022, Wang *et al*., 2022), or be used to study aspects of CDI disease that are generally confounded by toxin-mediated effects, such as early colonization events.

In light of the possible applications of NTCD, and the relative paucity of well-characterized NTCD strains, we present here a detailed analysis of L-NTCD03, a clade 4 non-toxigenic *C. difficile* strain.

## Materials and Methods

### Bacterial strains and growth conditions

L-NTCD03 was isolated from fecal material of a healthy elderly subject negative for HIV-1, HIV-2, HCV, HBV, as follows. Fecal material was stored in Stool Transport and Recovery (STAR) buffer and stored at −80C. Ethanol treated material was inoculated into *C. difficile* enrichment modified broth (CDEB MOB; Mediaproducts NV, Groningen, The Netherlands, cat no. 21.1295). After 6 days of growth at 37C under anaerobic conditions, the culture was plated onto *C. difficile* selective growth medium (CLO; VWR International, cat.no. 43431). From this medium, a colony was streaked to singles on a Tryptic Soy Sheep Blood medium (TSS; bioMérieux, cat.no. 43009,), and growth was harvested into glycerol-containing broth (Tritium Microbiology, cat. no. I200.36.0001) for storage at −80⁰C.

Control strains belonging to PCR ribotype 027 (LL-027) and 078 (LL-078) were obtained from the Leeds-Leiden collection (Knetsch *et al*., 2012, Baktash *et al*., 2022).

Strains were cultured in simulated ileal effluent medium (SIEM)(Minekus *et al*., 1999, Wiese *et al*., 2022), Brain-Heart Infusion (BHI) broth, BHI supplemented with yeast extract (BHI-Y; BHI with 0.5 % yeast extract) or CDMN (Aspinall & Hutchinson, 1992) medium supplemented with 0.1% sodium taurocholate (CDMNT).

### Molecular biology

A genomic region including *cfr*(B) and its native promoter was amplified by PCR using Q5 polymerase (NEB) and primers oIMM-01 and oIMM-02. The resulting PCR product was digested with KpnI and BamHI and cloned into similarly digested pAP24 (Oliveira Paiva *et al*., 2016). The resulting plasmid, pIMM03, was sequence verified using primers NF793, NF794 (Fagan & Fairweather, 2011), oIMM-01, oIMM02, oIMM-03 and oIMM-04 was introduced using standard laboratory methods into strain 630Δ*erm* (van Eijk *et al*., 2015, Roseboom *et al*., 2023). Transconjugants were verified by PCR using primers oWKS-1070/oWKS-1071 (targeting chromosomal *gluD*), oWKS-1387 and oWKS-1388 (targeting plasmid-located *traJ*) and oIMM-01/oIMM-02 (targeting *cfr*(B)) using appropriate controls. All oligonucleotide sequences are listed in **Table 3**.

### Antimicrobial susceptibility testing

To determine minimal inhibitory concentrations (MIC) of a panel of antibiotics epsilometer tests (Etests) were used. This panel consisted of cotrimoxazol, linezolid, metronidazole (Liofilchem), augmentin, cefuroxime, ciprofloxacin, chloramphenicol, clindamycin, erythromycin, meropenem, moxifloxacin, penicillin, rifampicin, tetracyclin, teicoplanin and vancomycin (bioMerieux). For this test, bacterial suspensions corresponding to a turbidity of 1.0 McFarland unit were plated on Brucella blood agar supplemented with 5 mg L^−1^ hemin and 1 mg L^−1^ vitamin K (bioMerieux) plates, unless indicated otherwise. MIC values were determined after 48 h of anaerobic incubation at 37⁰C as recommended by Clinical and Laboratory Standards Institute (CLSI), including relevant control strains (CLSI, 2012). Fidaxomicin MIC was determined using the agar dilution method according to CLSI guidelines on fresh Brucella Blood Agar supplemented with 5 mg L^−1^ hemin and 1 mg L^−1^ vitamin K. Epidemiological cut-off values were used as defined by European Committee on Antimicrobial Susceptibility Testing (EUCAST, 2026), when available.

MIC measurements for engineered strains (3 independent transconjugants) were performed on BHI-yeast extract agar with either an E-test (clindamycin, linezolid) (bioMerieux or Liophilchem, respectively) or by incorporating retapamulin (Sigma CDS023386) or streptogramin A/virginiamycin M1 (Sigma V2753) at 0-128 mg/L into the agar.

### PCR ribotyping

Capillary gel electrophoresis ribotyping of L-NTCD03 was performed according to standardized protocols (Fawley *et al*., 2015). In short, 2µl of isolated DNA was added to 23 µl of the master mix which includes the 16S and 23S primers and HotStart Taq Master Mix (Qiagen). Using a MyCycler thermocycler (Biorad), the resulting DNA mix was subjected to an enzyme activation step at 95 °C for 15 minutes followed by 24 cycles of amplification at 95 °C for 1 minute, 1 minute at 57 °C and 1 minute at 72 °C. Finally, the DNA mix was subjected to 72 °C for 30 minutes. The sample was prepared for analysis by adding PCR product as 1 µl of PCR product to 8.5 µl HiDi formamide (Applied Biosystems) and 0.5 µl of GeneScan™ 1200 LIZ marker (Thermo Fisher) and samples was denatured at 95 °C for 5 minutes. To determine the ribotype, it was then analyzed using the ABI3500XL (capillary length 50 cm) and the resulting *.fsa files were compared to a reference database in BioNumerics (bioMérieux).

### Toxin ELISA

To determine the productions of Toxin A and B, the *C. difficile* Tox A/B II^TM^ kit (Techlab) was used according to manufacturer’s protocol with a minor modification. In short, the supernatants that were collected at 0, 24, 48, 72 and 96 h from growth curves in SIEM pH=5.8 and pH=6.8 were filtered using a UNIFILTER microplate device (0.45 µm cellulose acetate filter, long drip, 96 wells, GE Health Care Life Sciences). The samples were applied to the filter plate and centrifuged at 3220 G for 5 minutes at 4°C. The filtrates were used in the analysis as described by manufacturer and the optical density was measured at 450 nm (OD_450_). A threshold value of OD_450_ of 0.12 was used determine whether toxins were present in the filtrate.

### Cell lines and culture conditions

Human colorectal adenocarcinoma cells (Caco-2) were obtained from the American Type Culture Collection (ATCC-HTB-37) and cultured in minimum essential medium (MEM, Gibco) supplemented with 10% bovine fetal serum (10% FCS), 1% non-essential amino acids (NEAA, Lonza), 2 mM glutamine (Sigma Aldrich), 100 units/ml penicillin (Sigma Aldrich), and 100 units/ml streptomycin (Sigma Aldrich). Epithelial-like, mucin-producing, LS174T cells were obtained from Sigma Aldrich (87060401) and cultured in the medium described for Caco-2. Both cells were grown to 80% confluency before experimental conditions were prepared.

### *In vitro* intestinal epithelial barrier model

The apical side of Thincert 12 well cell culture inserts (PET, pore size 0.4 µm, Greiner Bio-one) were coated with 250 µg/well purified human collagen type IV (OptiCol collagen Type IV, dissolved in 0.25% acetic acid, Cell Guidance Systems) and washed with sterile PBS after 1-2 h incubation.

To mimic the human colon environment, an 80:20 ratio mixture of Caco-2 and LS174T in MEM supplemented with 10% FCS, 1% non-NEAA, 2 mM glutamine, 100 U/ml penicillin, and 100 U/ml streptomycin (cell culture medium) was prepared. To function as controls, 100:0 and 0:100 ratio mixtures of Caco-2 and LS174T were prepared in cell culture medium. For all conditions, 1*10^5^ cells were seeded on the apical side of the PET membrane. To mimic the intestinal environment, medium was present in both the basal compartment (the well) and apical compartment (on top of the PET membrane) of the system. The systems were incubated at 37°C and 5% CO2. The cells in this model were allowed to grow and differentiate for 21 days during which the medium in both compartments was refreshed every other day.

To monitor the development of the *in vitro* intestinal epithelial barrier model, starting at T=0 the transepithelial electrical resistance (TEER) values were measured using EVOM-2 resistance meter (World Precision Instruments) every 7 days.

On day 21, the medium in the apical compartment of the system was replaced with a 10 times diluted, in cell culture medium, supernatant harvested at 72 h in a growth curve of all of the strains in BHI-y medium. This supernatant was incubated in the apical compartment for 1 hour at 37°C and 5% CO2, removed afterwards and replaced with cell culture medium. In every condition, a control well was exposed to calcium free medium (SMEM, Gibco) to determine as a positive control. TEER measurements were performed at 4 (only SMEM condition), 24, and 48 h post-exposure. Morphology was assessed under light microscopy every 24 hrs.

### Swine IsoLoop model

We compared the effect of L-NTCD03 and LL-078 on intestinal mucosa *in vivo* in the pig IsoLoop model, a translational surgical *in situ* ileal loop platform for host-microbiome-pathogen interface (Bayne *et al*., 2025). 12 h before the start of the surgery, 1 ml was drawn from a culture in the exponential growth phase of either L-NTCD-03 or LL-078 in BHI-y and inoculated into fresh, pre-reduced CDMNT medium and grown anaerobically at 37 °C. 10 ml of culture was aliquoted per loop and stored anaerobically at room temperature until injection was required.

3-month-old, female pigs (*Sus scrofa*, mixed breed, 50-60kg) were used under USDA pain category D, per Institutional Animal Care and Use Committee (IACUC) protocol IACUC-22-066 essentially as described (Bayne *et al*., 2025).

In short, the pig was placed in dorsal recumbency under general anesthesia. Ileal loops were rinsed with sterile saline solution before treatment with an antibiotic cocktail (60 mg/L norfloxacin; 160 mg/L moxalactam) for 30 minutes. After a saline rinse, each independent loop except for the most aboral segment was inoculated using a tuberculin needle full thickness into the lumen. Upon recovery, animals were housed under standard husbandry conditions with regular veterinary assessments to ensure welfare. The loops were sampled at euthanasia by cutting the mesentery and immediately cooling the intestines on ice. On ice, the loops were cut open one by one and a piece of tissue, including intestinal content, was fixed in 10% neutral buffered formalin (VWR) before embedding in paraffin.

5μm-thick tissue sections were stained with hematoxylin and eosin. Microscopic analysis of histopathologic features was conducted by a board-certified pathologist, blinded to sample identities. The scoring rubric, including descriptive statistics applied, is provided as **Supplementary Information**, and raw data of the scoring is included as **Supplementary Table 1**. Scores are reported as total histological score per ileal loop (range 0-20), computed as the sum for four 0-5 ordinal components (epithelial necrosis, mucosal/lamina propria edema, crypt abscess and neutrophils in villi/lamina propria). Statistical analyses of histological scores were performed in Python (NumPy/SciPy); figures were generated with matplotlib. All tests were two sided with α = 0.05.

### DNA extraction and genome sequencing

To extract pure high-molecular weight total DNA, a two-fold serial dilution of L-NTCD03 in BHI-Y medium was prepared. After overnight incubation at 37⁰C in an anaerobic environment, 4mL of culture with an optical density at 600nM between 0.6 and 0.8 was spun down and DNA was extracted from the cell pellet using the 500/G genomic tip kit (Qiagen), according to the instructions of the manufacturer. Quantity and quality of the DNA was verified by gel electrophoresis on a 0.8% agarose gel, using the Qubit dsDNA high-sensitivity kit (Qubit) and a Femto Pulse system (Agilent), according to the manufacturer’s protocols.

PacBio sequencing libraries were generated according to the manufacturer’s multiplexed microbial library preparation protocol, part number 101-696-100, version 7, July 2020 release (Pacific Biosciences) using the SMRTbell Express Template Prep Kit v2.0 with the following modifications: genomic DNA was sheared using Speed 34 on the Megaruptor 3 (Diagenode) and an additional size selection step of 6-50kb fragments on the Blue Pippin (Sage Science) was included for the final SMRT bell library. The libraries were sequenced on a Sequel II platform (Pacific Biosciences) using the Sequel II Binding Kit v2.0, Sequencing Primer v4, Sequencing Kit v2.0 and a 30hr movie time.

### Genome annotation and bioinformatics analyses

De novo genome assembly was carried out using Hifiasm (version 0.16.1-r375; parameters ‘--hg-size 4m’) (Cheng *et al*., 2021) and the start position was set to *dnaA* using Circlator fixstart (version 1.5.5 using default parameters) (Hunt *et al*., 2015). Completeness check was performed using Busco (version 5.2.2; parameters ‘-m genome -l clostridia_odb10’) (Manni *et al*., 2021) and assembly statistics were calculated and visualised with Quast (version 5.0.2; using NZ_CP076401 as reference genome)(Gurevich *et al*., 2013). Finally, Prokka (version 1.14.6; parameters ‘--genus Clostridioides --species difficile --compliant’)(Seemann, 2014) was used to annotate the genome.

A database of AMR genes was constructed using the AMRFinder plus tool v3.10.23 (parameters “-n” “–organism Clostridioides difficile”, “—plus”) to identify antimicrobial resistance (AMR) genes (Feldgarden *et al*., 2021). Minimal “% coverage of reference sequence” and “% Identity to refence sequence” were set to 60% and 90% percent, respectively.

BacAnt (Hua *et al*., 2021) and MobileElementFinder (Johansson *et al*., 2021) were used to identify putative transposable elements. progressiveMauve was used to align genome sequences (Darling *et al*., 2010).

Multi-locus sequence typing (MLST) was performed using mlst (v2.19.0) with the PubMLST *C. difficile* database updated to October 21, 2021, (Jolley *et al*., 2018) and verified on December 1, 2025.

Genomic comparisons were made in Geneious R10 (BioMatters Ltd), and figures for publication were prepared using CAGECAT (van den Belt *et al*., 2023) and Adobe Illustrator 26.3.1.

### Data availability

L-NTCD03 genome information is available under BioProject PRJEB62683 (raw reads: ERR11495271, assembly: ERZ18456945). The annotated genome is OX637968.1.

## Results

### L-NTCD03 belongs to a rare PCR ribotype, RT416

As part of a study investigating carriage of multidrug resistant organisms and *C. difficile* in nursing homes (Ducarmon *et al*., 2021), fecal material was obtained from a healthy elderly subject. *C. difficile* was cultured from this material as described in Materials and Methods. The resulting isolate was negative in a multiplex PCR that targets the toxin genes *tcdA*, *tcdB, cdtA* and *cdtB* (Control, 2018), suggesting that the isolate was non-toxigenic. Hereafter, the isolate is referred to as L-NTCD03.

We performed capillary electrophoresis PCR ribotyping (Fawley *et al*., 2015) of L-NTCD03 through the Dutch Expertise Center for *C. difficile*, hosted at the LUMC. We initially failed to obtain a ribotype assignment, as the banding pattern was not present in our database. The *.fsa files were shared with the *Clostridioides difficile* Ribotyping Network Laboratory (CDRN; PHE Microbiology Services, Leeds, UK), and they determined the isolate to belong to PCR ribotype 416. Since this identification, no other isolates were identified with the same ribotype in the Dutch National Expertise Center for *C. difficile* infections. This suggests a prevalence in the Netherlands that is below 0.01% on the basis of the number of strains typed since then.

### L-NTCD03 is susceptible to clinically used antimicrobials

To determine whether L-NTCD03 the minimal inhibitory concentration for different antimicrobials, we performed antimicrobial susceptibility testing using agar dilution (for fidaxomicin) or Etest (augmentin, cefuroxime, chloramphenicol, clindamycin, cotrimoxazol, erythromycin, linezolid, metronidazole, meropenem, moxifloxacin, penicillin, rifampicin, tetracyclin, teicoplanin and vancomycin) methods on Brucella Blood Agar (BBA), as recommended by CLSI (CLSI, 2012). Out of the 17 antimicrobials tested, L-NTCD03 demonstrated elevated MICs for cefuroxime, chloramphenicol, ciprofloxacin, erythromycin, linezolid, rifampicin and tetracycline (**Table 1**). In contrast, the strain is susceptible to antimicrobials that are used in CDI therapy - fidaxomicin, vancomycin and metronidazole – as well as several others (**Table 1**). As BBA is a suboptimal medium for determining resistance to metronidazole, we also retested the isolate on recently recommended Fastidious Anaerobe Agar supplemented with horse blood (Freeman *et al*., 2025); this confirmed susceptibility to metronidazole (data not shown). These results demonstrate that L-NTCD03 would likely be suppressed or eradicated using standard antimicrobial therapy for CDI, but not by certain other antimicrobials that increase susceptibility for CDI.

**Table 1.** Antimicrobial susceptibility of L-NTCD03.

| Antibiotic | MIC (mg/L)<sup>a</sup> | EUCAST (T)ECOFF (mg/L) | Putatively associated AMR gene<sup>b</sup> |
| --- | --- | --- | --- |
| Augmentin | 0.25 | - | - |
| Cefuroxime | >256 | - | *blaCDD* |
| Chloramphenicol | >256 | - | *catP*, *cfr*(B) |
| Ciprofloxacin | >32 | (32) | *gyrB_S366A* |
| Clindamycin | >256 | 16 | *cfr*(B), *cplR* |
| Cotrimoxazole | 0.06 | - | - |
| Erythromycin | 48 | 4 | *erm*(52) |
| Fidaxomicin | 0.06 | 0.5 | - |
| Linezolid | 6 | - | *cfr*(B) |
| Meropenem | 1 | (8) | - |
| Metronidazole | 0.06 | 2 | - |
| Moxifloxacin | 1 | 4 | - |
| Penicillin | 0.75 | - | *blaCDD* |
| Rifampicin | >32 | 0.004 | *rpoB_R505K* |
| Tetracycline | 2 | (0.25) | *tet*(M) |
| Teicoplanin | 1 | - | *vanZ1* |
| Vancomycin | 0.38 | 2 | - |

<sup>a</sup> MICs were determined using Etest, except for fidaxomicin (where agar dilution was used) as described in

### Supernatant from L-NTCD03 does not contain detectable toxin A and B

To confirm the absence of toxins, as suggested by the multiplex PCR result, we assessed the presence of TcdA and TcdB in culture supernatant using an enzyme-coupled immune assay (Techlab). We used a medium that mimicks the physiological conditions in the gut: simulated ileal effluent medium, SIEM (Minekus *et al*., 1999, Wiese *et al*., 2022).

During 96 h of growth in SIEM, toxin levels were below the limit of detection for all supernatants derived from L-NTCD03, as well as for the reference non-toxigenic strain LL-010 (**Figure 1A**). In contrast, toxins were readily detected in culture supernatants derived from the toxigenic reference strains LL-027 and LL-078 (**Figure 1A**).

**Figure 1.**
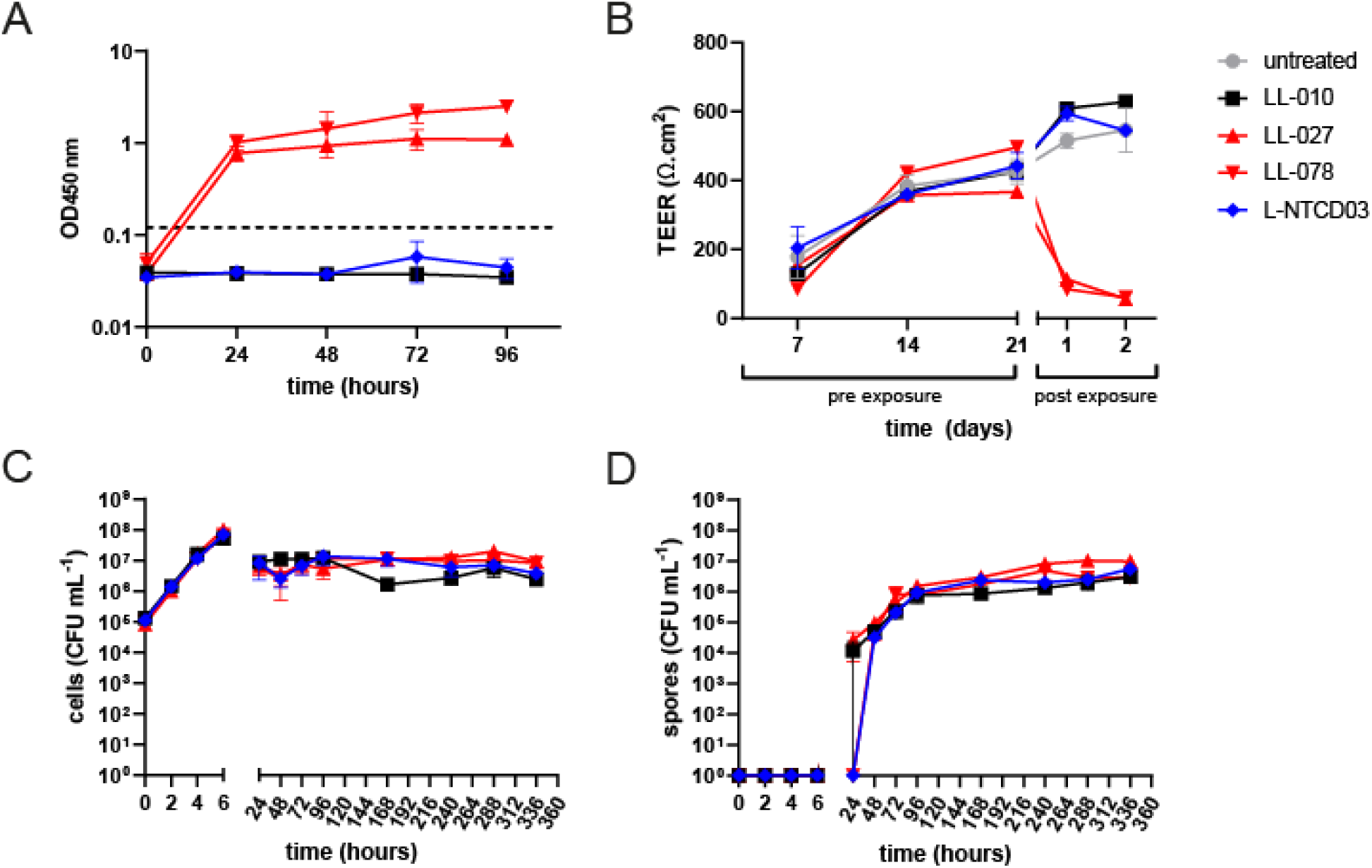
L-NTCD03 is a non-toxigenic *C. difficile* strain. **A.** L-NTCD03 does not show production of toxins in an ELISA assay. The dotted line indicates the threshold value indicated by the manufacturer for defining toxin production. **B**. Culture supernatant from L-NTCD03 does not lead to a reduction in transepithelial resistance as measured in a Caco2/LS174T cellular model that was matured for 21 days (pre-exposure) before applying culture supernatant (post exposure). **C**. Growth of L-NTCD03 is similar to other *C. difficile* strains. Colony forming units (CFU) per milliliter culture were determined by a droplet-plating method. **D**. Spore formation of L-NTCD03 is similar to other *C. difficile* strains. Spores were enumerated after ethanol treatment using a droplet plating method. Values indicated are the mean and standard error of the mean of n=3 biological replicates. LL-010 is a non-toxigenic control strain and LL-027 and LL-078 are toxigenic controls.

### Supernatant from L-NTCD03 does not affect transepithelial electrical resistance

Though our results clearly indicate the absence of TcdA and TcdB, we considered that there could be other factors that affect epithelial integrity, not detected in our assays so far. For this reason, we assessed the effect of culture supernatant derived from L-NTCD03, LL-010, LL-027 and LL-078 in an *in vitro* model for epithelial integrity. In this model, a mixture of Caco2 and mucin-producing LS174T cells is grown in a transwell system and the transepithelial electrical resistance (TEER), a proxy for epithelial integrity, is measured using electrodes.

The TEER values gradually increased over the first 14 days of incubation, consistent with cells growing to confluency, and stabilized between days 14 and 21 (**Figure 1B**). When filtered supernatants from 72h-cultures in BHI were applied to the cell layer, no drop in TEER was observed for the non-toxigenic strains (LL-10 and L-NTCD03), whereas LL-027 and LL-078 derived supernatants resulted in a significant decrease (**Figure 1B**). Similar results were obtained when only LS174T cells or Caco2 cells were used (data not shown). This shows that no other factors are produced by the non-toxigenic *C. difficile* strains that affect TEER in this cellular model.

### L-NTCD03 grows similar to other isolates in a simulated ileal effluent medium

To assess the growth and sporulation of L-NTCD03 in relation to well-characterized common subtypes of *C. difficile* in SIEM, we monitored the total number of colony forming units (CFU) per milliliter (mL) over a period of 14 days. Spore formation was monitored by enumeration of ethanol resistant CFUs.

Growth of L-NTCD03 was indistinguishable from LL-010, LL-027 and LL-078 for the first 6 h, during which the number of CFU/mL increased ∼3 log (**Figure 1C**). During stationary phase, we observed a lower total cell count of ∼10^7^ CFU/mL, for all strains (**Figure 1C**).

No spore production was found during the first 6h of growth in SIEM (**Figure 1D**). For LL-010 and LL-027 spores were detectable at 24h, but for L-NTCD03 and LL-078 spores were first found at the 48h-timepoint. Beyond this point, sporulation dynamics were comparable between all strains **Figure 1D**).

Thus, under the conditions tested, L-NTCD03 shows robust growth and sporulation, in line with other *C. difficile* strains.

### Phenotype microarrays show a common carbon utilization profile for L-NTCD03

In order to assess the carbon utilization by L-NTCD03 we performed phenotype microarrays on Biolog PM1 and PM2 plates, and analyzed the results using AMIGA software, as described previously (Midani *et al*., 2021). We used an arbitrary cut-off of 1.5-fold increase in or a 2-fold decrease in optical density to define differential growth. We observed stimulated growth on the carbon sources N-Acetyl-D-Glucosamine, D-Mannose, D-Mannitol, D-Fructose, alpha-D-Glucose, N-AcetylNeuraminic Acid, D-Melezitose, D-Tagatose and D-Glucosamine (**Supplemental Table S1**). A decreased cell density was observed in the presence of Glyoxylic Acid and Glycyl-LGlutamic Acid (**Supplemental Table S1**).

In parallel experiments, we observed similar profiles for representative strains from PCR ribotype 010, 027 and 078 (data not shown)(Collins *et al*., 2018, Midani *et al*., 2025). We conclude that L-NTCD03 carbon utilization is similar to that of other *C. difficile* isolates.

### L-NTCD03 does not induce *C. difficile*-associated intestinal lesions in the swine IsoLoop model

We have previously demonstrated the utility of the swine IsoLoop model for assessing *C. difficile*–associated lesions in a translationally relevant gut microbiota–pathogen interface (Bayne *et al*., 2025). Using this platform, we compared the effects of L-NTCD03 in relation to the effects induced by the toxigenic reference strain LL-078.

We inoculated multiple loops in the same animal (one animal per strain, either LL-078 or L-NTCD03), and compared the histology to control loops without *C. difficile* and normal ileum (**Figure 2)**. Loops inoculated with LL-078 showed signs of inflammation, whereas those inoculated with L-NTCD03 were comparable to the control loop (**Figure 2A**). The total histological score (further detailed in the Methods section of this manuscript) of samples derived from loops inoculated with LL-078 were significantly higher than those derived from, whereas the histology of L-NTCD03 derived samples was comparable to that of control loops (**Figure 2B** and **Supplemental Table 1**).

**Figure 2.**
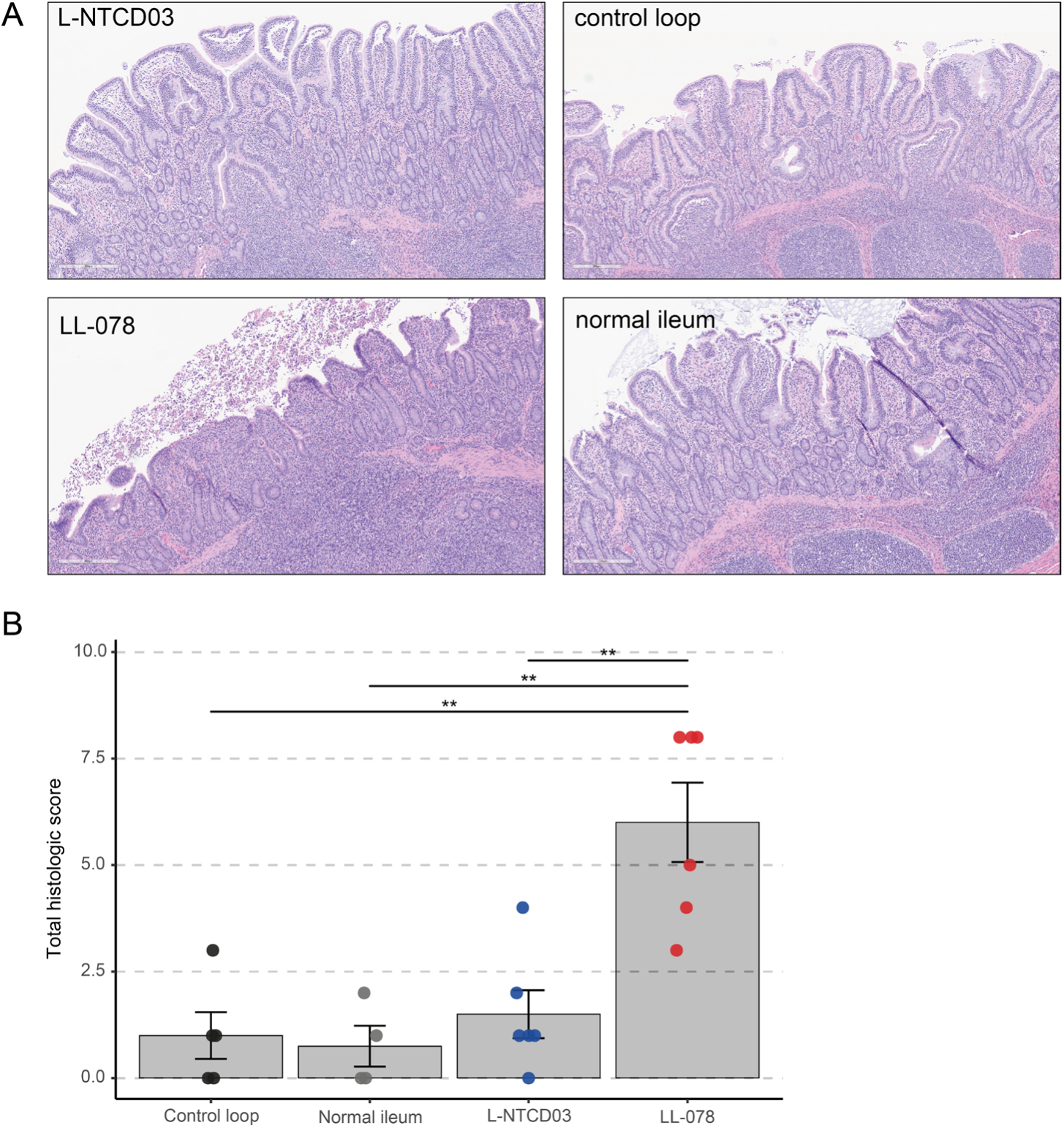
L-NTCD03 does not induce pathology in the swine IsoLoop model. **A.** Representative images from histological analyses. Tissues samples were formalin fixed and paraffin embedded, sectioned and hematoxylin and eosin stained. No significant *C. difficile*-associated lesions are observed in the control loop (an ileal loop that was antibiotic-treated but not inoculated with *C. difficile*), normal ileum (not a loop but control for surgical integrity of the loops) and the loop inoculated with L-NTCD03. In contrast, the loop inoculated with LL-078 (toxigenic strain, positive control for *C. difficile* pathology) showed blunting of the villi with epithelial necrosis, neutrophilic infiltration of lamina propria, and crypt abscesses. **B**. Combined histological scores for the different samples. A bar plot depicting the mean histological score with standard error of the mean is given, with individual datapoints depicted. Prespecified contrasts (see Supplementary Information) are shown with brackets, with asterisks indicating the raw p-value magnitude, for readability (**: p < 0.01), while significance is determined by Holm adjustment across the three prespecified tests.

### L-NTCD03 is an ST39 (clade 4) isolate carrying a *cfr*(B) resistance gene

*C. difficile* is a phylogenetically diverse species, with isolates falling into at least 5 classical (CDI-associated) clades and 3 so called cryptic clades, that have been proposed to form separate genomospecies (Knight *et al*., 2021). To place the L-NTCD03 in phylogenetic context, we reconstructed the complete genome sequence of the strain using Pacific Biosciences circular consensus sequencing (BioProject PRJEB62683). After assembly, the resulting genome showed a single circular contig of 4,318,385 bp – indicating the absence of extrachromosomal elements - and an average [G+C]-content of 28,91%. Reference assembly of short-read (Illumina) sequence data did not identify any nucleotide variants (data not shown), underscoring the quality of the assembly.

We derived the 7-gene multilocus sequence type of our isolate using PubMLST (Griffiths *et al*., 2010, Jolley *et al*., 2018). L-NTCD03 was found to belong to ST39 (*adk* 3, *atpA* 7, *dxr* 10, *glyA* 8, *recA* 7, *sodA* 2, *tpi* 10), which falls into clade 4. A core genome phylogenetic tree of representative isolates of different *C. difficile* clades using Panaroo and IQTree confirmed this placement (data not shown)(Minh *et al*., 2020, Tonkin-Hill *et al*., 2020, Baktash *et al*., 2022).

We performed automated annotation of the genome sequence using Prokka (Seemann, 2014). This resulted in 3,844 predicted coding sequences, 88 tRNAs, 34 rRNAs, 1 tmRNA and 8 repeat regions. As expected, no complete genes corresponding to the toxin genes *tcdA*, *tcdB*, *cdtA* and *cdtB* were detected; instead the PaLoc insertion site conforms to what has been described for clade 4 strains (115bp insertion)(Dingle *et al*., 2014).

We used AMRfinderPlus to identify antimicrobial resistance genes in the L-NTCD03 genome (Feldgarden *et al*., 2022). We found 14 putative antimicrobial resistance genes, which are associated with 12 different classes of antibiotics (**Supplemental Table 2**); ten of these might be linked to the phenotypically observed reduced susceptibility/resistance (**Table 1**).

Though *cfr*(B) has been implicated in antimicrobial resistance of *C. difficile* to PhLOPSa (phenicol, lincosamide, oxaxolidinone, pleuromutilin and streptogramin A) antimicrobials (Marin *et al*., 2015, Stojkovic *et al*., 2019), a direct causal role for this gene in resistance to these antimicrobials has not been shown to date. Therefore, we cloned the *cfr*(B) gene under its native promoter in a shuttle plasmid and introduced this plasmid into the commonly used *C. difficile* laboratory strain 630Δ*erm* (Hussain, 2005, van Eijk *et al*., 2015). We assessed antimicrobial susceptibility using Etest and found that introduction of *cfr*(B) increased the minimal inhibitory concentration for clindamycin, linezolid, retapamulin and streptogramin A (**Table 2**) in comparison to a control strain carrying a similar plasmid that carried the reporter gene *sluc*^opt^ (Oliveira Paiva *et al*., 2016) rather than *cfr*(B). The MIC increased >128-fold for clindamycin (from 2 mg/L to >256 mg/L), 12-fold for linezolid (from 1 mg/L to 12 mg/L), 64-fold for retapamulin (from 0.125 mg/L to 16 mg/L) and 8-fold for streptogramin A (from 2 mg/L to 16 mg/L) in the presence of *cfr*(B). We observed no change in MIC for vancomycin and metronidazole (data not shown). This result indicates that *cfr*(B) can indeed confer resistance to lincosamide, oxazolidinone, pleuromutilin and streptogramin antimicrobials in *C. difficile*, consistent with expectations (Marin *et al*., 2015, Stojkovic *et al*., 2019).

**Table 2.** Table 1. Minimal inhibitory concentrations of *C. difficile* 630Δ*erm*-derived strains carrying different plasmids.

| plasmid | antimicrobial MIC (mg/L) |  |  |  |
| --- | --- | --- | --- | --- |
|  | clindamycin | linezolid | retapamulin | streptogramin A |
| pAP24 (control) | 2 | 1 | 0.125 | 2 |
| pIMM03 (*cfr*(B)) | >256 | 12 | 16 | 16 |

Experiments were performed at n>3, with results consistent between experiments. Clindamycin and linezolid MICs were determined by Etest, and MICs for retapamulin and streptogramin A were determined by modified agar dilution (see Methods).

**Table 3.**
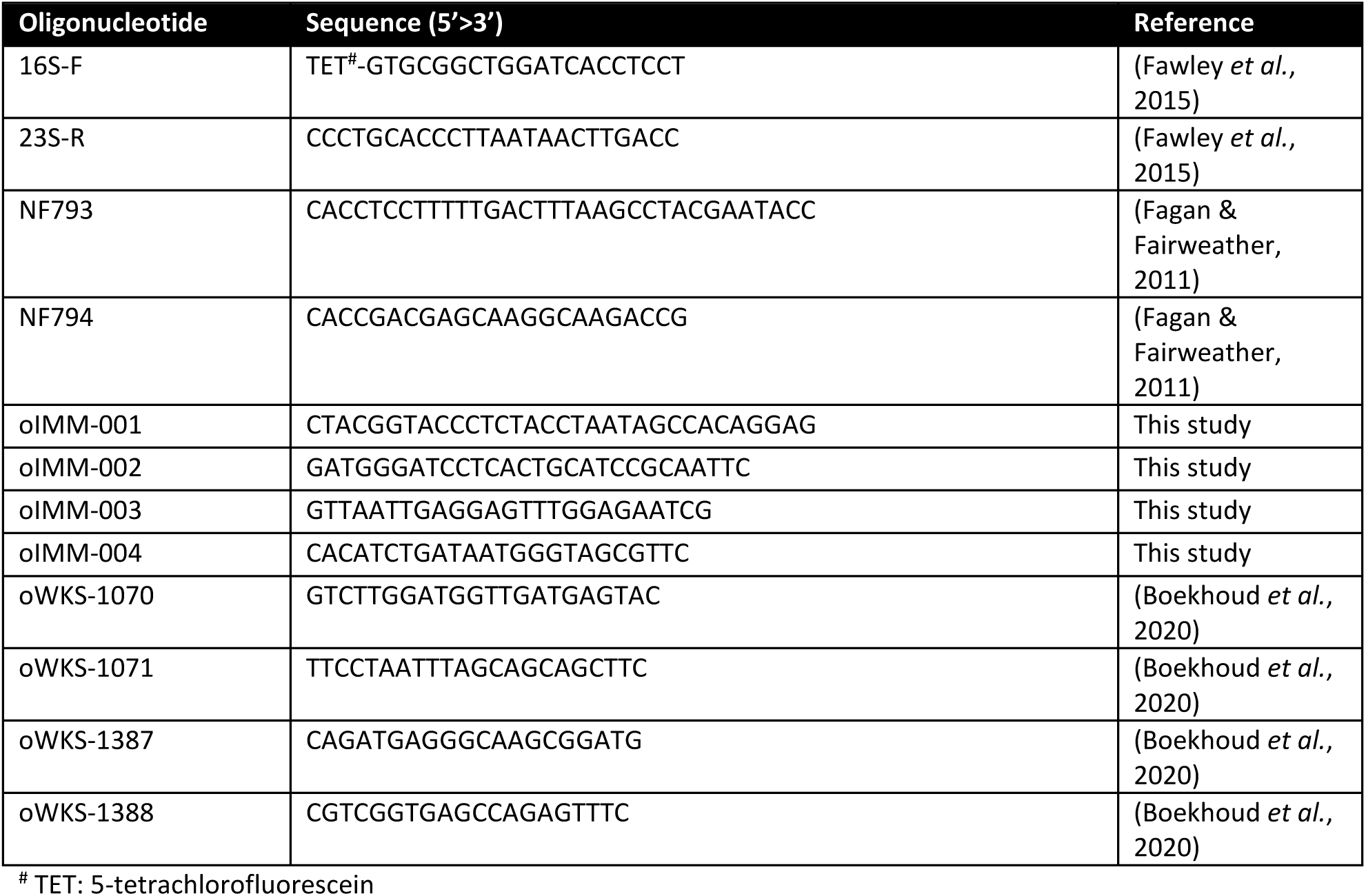
Oligonucleotides.

### The genome of L-NTCD03 harbors several putative mobile genetic elements

The BacAnt annotation server (Hua *et al*., 2021) identified two putative transposable elements, with homology to Tn*4451* (GenBank U15027; 94.57% identity and 100% coverage; bp 524023-530368) and Tn*5405* (GenBank U73027; 99.84% identity and 95.97% coverage; bp 551080-553654). Of note, the *catP* gene is contained on the former, suggesting a potential for transmission of this resistance determinant. MobileElementFinder (Johansson *et al*., 2021), in addition to Tn*4551*, identifies a chromosomal region with homology to Tn*6009* (GenBank EU399632; identity 99.95% and coverage 100%; bp 268640-270528). Comparison with a closely related clade 4 genome (DSM 29629, GenBank NZ_CP016104) reveals that this is in fact a 17.9-kb transposon highly similar to Tn*916*, which has inserted in a gene (*pdeR*) encoding a GGDEF domain containing protein (in L-NTCD03 this gene is split into two ORFs: LLHAOGBK_00311 and LLHAOGBK_00328) (**Figure 3**). The transposon (**GenBank OR192611**) is also found in several *Streptococcus* and *Enterococcus* genome assemblies (data not shown), and was assigned the number Tn*7696* by the transposon registry (Tansirichaiya *et al*., 2019). Compared to Tn*916*, Tn*7696* has a 120-bp in-frame deletion in the gene encoding a putative endopeptidase p60 precursor (ORF14 in Tn*916*, LLHAOGBK_00319 in L-NTCD03) as well as several SNPs.

**Figure 3.**
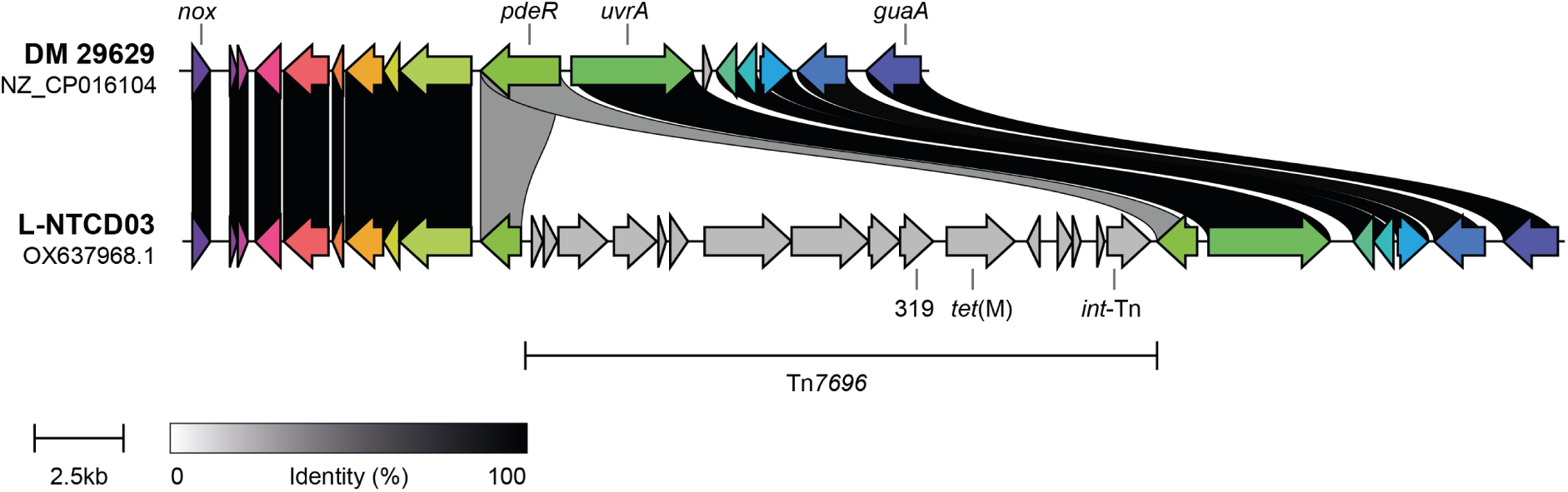
L-NTCD03 harbors the novel transposon Tn*7696*. Comparison of a region of the genome of L-NTCD03 with the genome of the closely related clade 4 strain DSM 29629 (GenBank NZ_CP016104). The insertion of Tn*7696* into the *pdeR* gene (LLHAOGBK_00328 and LLHAOGBK_00311/CDIF29629_RS01565) is evident. Tn*7696* harbors the tetracycline resistance gene *tet*(M) and an integrase (*int-*Tn), like Tn*916*. The gene with locus tag LLHAOGBK_00319 (see main text) is abbreviated to 319 for simplicity. Colors and linkages indicate homology between genes. *nox* encodes a predicted NADH dehydrogenase/nitroreductase (LLHAOGBK_00337/ CDIF29629_RS01610), *uvrA* an excinuclease (LLHAOGBK_00310/CDIF29629_RS01560) and *guaA* a glutamine-hydrolyzing GMP synthase (LLHAOGBK_00305/CDIF29629_RS01525).

Finally, we employed two tools to predict i) putative phages (Wishart *et al*., 2023) and ii) genomic islands (Bertelli *et al*., 2017). This identified two possible complete (pro)phages in L-NTCD03, with homology to phiCDHM19 (NC_028996; bp 664975-719994) and phiC2 (NC_09231; bps 2626715-2687566) both of which are also predicted with high confidence by geNomad (Camargo *et al*., 2024), and 2 large (bp 490582-569665 and 3900143-3992891) plus several small regions that are likely horizontally acquired.

Overall, the complete genome sequence suggests a clear relation with other clade 4 *C. difficile* isolates, and a significant contribution of horizontal gene transfer to its genetic content.

## Discussion

Here, we provided a characterization of L-NTCD03, isolated from a healthy elderly subject. We confirmed that this isolate does not express the large clostridial toxins *tcdA* and *tcdB*, using ELISA, and showed that supernatants derived from an L-NTCD03 culture do not induce a loss of TEER in a trans-well model of colonic epithelium. Additionally, data from the swine IsoLoop model suggest that there is no induced pathology in loops inoculated with L-NTCD03 compared to controls. The loop experiment provides insight into strain behavior within a translationally relevant, *C. difficile*–predisposing intestinal environment, using a customized dysbiotic gut segment that enables precise local control of the bacterial inoculum, bile acid milieu, and other key microenvironmental parameters. The data is in line with the genomic analysis, which shows a lack of identifiable toxin genes, including the binary toxin locus. Together, these data strongly suggest that L-NTCD03 is unlikely to induce disease.

The high-quality complete genome generated for L-NTCD03 as part of this work, allowed us to place the PCR ribotype 416 strain in clade 4. The most closely related publicly available genome at the time of analysis is DSM 29629 (Riedel *et al*., 2017)(CP016104.1). Like L-NTCD03, this strain was non-toxigenic and assigned to clade 4, however, it is listed as ribotype SLO 235. At present it is unknown if the ribotyping patterns for SLO 235 and RT416 (as determined by capillary ribotyping and assigned at the CDRN) are similar or the same. Interestingly, in the manuscript describing DSM 29629 the authors note metabolic versatility between 17 different *C. difficile* strains that does not appear to necessarily follow the 5 major phylogenetic clades (clade) or PCR ribotypes tested. In the present work, we find that L-NTCD03 demonstrated a carbon utilization profile that is quite similar to that of other *C. difficile* strains under the conditions tested. We have to note that this was assessed at a single concentration and we cannot rule out that differences may exists at alternative concentrations, similar to what has been noted for other *C. difficile* strains belonging to other ribotypes (Collins *et al*., 2018, Midani *et al*., 2025).

We have determined the phenotypic antimicrobial resistance profile for 17 antimicrobials and found that several could be explained by genetic determinants identified in the complete genome of L-NTCD03. Some of these determinants appear to be associated with mobile genetic elements (Bannam *et al*., 1995, Derbise *et al*., 1997, Soge *et al*., 2008, Hua *et al*., 2021), supporting the view that NTCD strains may act as a reservoir for resistance determinants (Imwattana *et al*., 2020). As observed also by others for NTCD strains (Imwattana *et al*., 2020), we also observed resistance to rifampicin and ciprofloxacin in L-NTCD03. It is interesting to speculate that high levels of resistance in NTCD may relate to prolonged carriage, or reflect antibiotic use in geographic areas where these strains are common. Rifampicin, for instance, is commonly used in the treatment of tuberculosis patients. For clade 4 strains, high prevalence in Asia has been noted (Imwattana *et al*., 2020) but such strains are also more broadly found (Viprey *et al*., 2022). Notably, though the *nimB*-Cd resistance gene was identified (**Supplemental Table 2**), we observed no resistance to metronidazole. Consistent with this, *nimB* is commonly found in susceptible isolates of *C. difficile* and L-NTCD03 lacks the P*nimB*^G^ mutation associated with increased *nimB* expression and inducible metronidazole resistance (Olaitan *et al*., 2023). It also highlights the putative nature of automated identification of antimicrobial resistance determinants.

Finally, we identified the *cfr*(B) gene in L-NTCD03. The *cfrB* gene encodes a rRNA methyltransferase that confers multidrug resistance by ribosomal target modification and, in addition to *C. difficile*, has been found in *Enterococcus* spp and methicillin-resistant *Staphylococcus aureus*. We showed that this gene can confer resistance to clindamycin (>256 mg/L), linezolid (12 mg/L), repatamulin (16 mg/L) and streptogramin A (16 mg/L) in the RT012 laboratory strain 630Δ*erm* (van Eijk *et al*., 2015). Our data confirm that *cfr*(B) is a legitimate PhLOPSa (phenicol, lincosamide, oxazolidinone, pleuromutilin, streptogramin A) resistance gene. We were unable to establish whether *cfr*(B) plays a role in phenicol resistance, though, as L-NTCD03 also contains the *catP* (chloramphenicol resistance gene) and the plasmid we used to introduce the *cfr*(B) gene in our laboratory strain also carries a *catP* resistance determinant (Fagan & Fairweather, 2011, Oliveira Paiva *et al*., 2016). We have, however, independently observed that *catP*-negative clinical isolates that carry *cfr*(B) demonstrate elevated chloramphenicol MICs compared to isolates that lack *cfr*(B), suggesting that *cfr*(B) also affects chloramphenicol MICs. Though it has previously been shown that *cfr* gene carriage can be associated with linezolid resistance in *C. difficile* (Marin *et al*., 2015, Stojkovic *et al*., 2019) or lead to an increase in MIC for certain antibiotics when expressed in a heterologous host (Hansen & Vester, 2015), to our knowledge, the present study is the first direct demonstration of this function for *cfr*(B) as a PhLOPSa resistance gene in this organism.

## Acknowledgements

We thank members of our laboratory as well as prof. dr. A.P. Roberts and prof. dr. L.C. Fortier for helpful discussions. We are indebted to prof. dr. P. Hiemstra and his team at LUMC for support with the TEER measurements and the CDRN (prof. M.H. Wilcox and dr. W. Fawley) for the ribotype assignment for this strain.

## Author contributions

BN, EJK and WKS conceived the study. CH and IMJGS performed typing and susceptibility testing, BN performed characterization of L-NTCD03 in *in vitro* and cellular assays. RHAMV and SLK performed PacBio sequencing. SN and QRD performed bioinformatics analyses. CKB and RAB performed the Biolog experiments. JB, CC, MLM, AL, BS, AR and SM conducted the animal experiments. FY performed the histological scoring. EB, SM and TO reviewed the histology. WKS, TO, SM and EJK supervised work. BN and WKS drafted the manuscript. All authors edited the manuscript and agreed to its submission.

## Supplementary information

### Histological scoring

H&E-stained sections from each ileal loop were graded for four components present in the dataset: epithelial necrosis, edema (mucosal/lamina propria), crypt abscess, and neutrophils in villi/lamina propria. A fifth field, submucosal edema, was also recorded but was invariant (0 in this experiment) and therefore excluded from totals.

Each component used an ordinal 0–5 scale with shared anchors; the best matching description was assigned:

- Epithelial necrosis: 0 = absent; 1–2 = scattered villus-tip necrosis; 3 = multifocal epithelial lifting/patchy denudation; 4–5 = broad denudation/ulceration with diffuse surface loss.
- Edema (mucosal/lamina propria): 0 = none; 1–2 = subtle to mild expansion; 3 = clear expansion with villus blunting; 4 = marked expansion; 5 = diffuse expansion with severe blunting/separation.
- Crypt abscess: 0 = none; 1–2 = rare crypts with a few intraluminal neutrophils/debris; 3 = several crypts with clear pus; 4–5 = numerous/widespread abscesses across fields.
- Neutrophils in villi/lamina propria: 0 = no neutrophils; 1= scattered individual cells; 2 = multifocal small clusters; 3 = multifocal large clusters; 4 = confluent clusters; 5 = sheets/near-diffuse infiltration. (Field counts may aid consistency; final grade is by pattern/severity.)

Total histologic score: per loop, the sum of the four variable components (range 0–20); higher totals indicate greater injury. Each loop was treated as the experimental unit; scoring was performed blinded to group identity.

### Statistical analyses

#### Outcome and unit

The primary endpoint was the total histologic score per ileal loop (range 0–20), computed as the sum of four 0–5 ordinal components: epithelial necrosis, mucosal/lamina propria edema, crypt abscess, and neutrophils in villi/lamina propria. Submucosal edema values were invariant and excluded from totals. Each loop was the experimental unit; scoring was blinded to group.

#### Descriptive statistics

Group data are summarized as median [Q1–Q3]. Figures display mean ± SEM with per loop jittered points.

#### Global comparison

Overall heterogeneity among Control loop, Normal Ileum, L-NTCD03 and LL-078 was assessed with the Kruskal–Wallis test (two sided, α = 0.05) using mid ranks with tie correction.

#### Primary prespecified contrasts

The biological comparisons of primary interest were LL-078 vs Control loop, LL-078 vs Normal ileum, and LL-078 vs L-NTCD03. Each was tested using a two sided Mann–Whitney U test (Wilcoxon rank sum). Multiplicity was controlled across these three tests using the Holm procedure (family wise α = 0.05). For each contrast we report the Holm adjusted p value and the rank biserial correlation (r) as an effect size.

**Supplementary Table 1.**
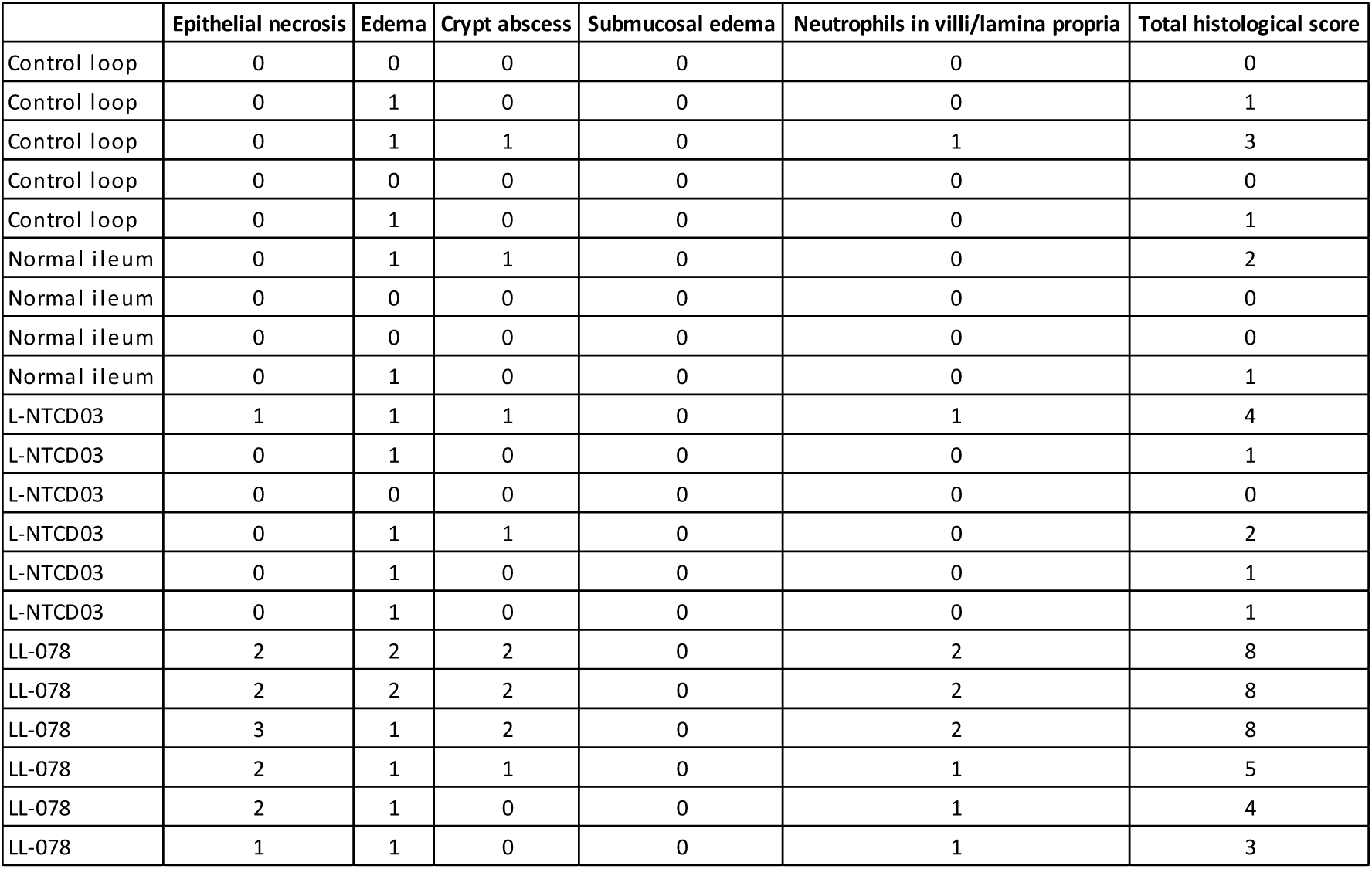
Histopathological analysis of the pig-ileal loop experiments.

|  | Epithelial necrosis | Edema | Crypt abscess | Submucosal edema | Neutrophils in villi/lamina propria | Total histological score |
| --- | --- | --- | --- | --- | --- | --- |
| Control loop | 0 | 0 | 0 | 0 | 0 | 0 |
| Control loop | 0 | 1 | 0 | 0 | 0 | 1 |
| Control loop | 0 | 1 | 1 | 0 | 1 | 3 |
| Control loop | 0 | 0 | 0 | 0 | 0 | 0 |
| Control loop | 0 | 1 | 0 | 0 | 0 | 1 |
| Normal ileum | 0 | 1 | 1 | 0 | 0 | 2 |
| Normal ileum | 0 | 0 | 0 | 0 | 0 | 0 |
| Normal ileum | 0 | 0 | 0 | 0 | 0 | 0 |
| Normal ileum | 0 | 1 | 0 | 0 | 0 | 1 |
| L-NTCD03 | 1 | 1 | 1 | 0 | 1 | 4 |
| L-NTCD03 | 0 | 1 | 0 | 0 | 0 | 1 |
| L-NTCD03 | 0 | 0 | 0 | 0 | 0 | 0 |
| L-NTCD03 | 0 | 1 | 1 | 0 | 0 | 2 |
| L-NTCD03 | 0 | 1 | 0 | 0 | 0 | 1 |
| L-NTCD03 | 0 | 1 | 0 | 0 | 0 | 1 |
| LL-078 | 2 | 2 | 2 | 0 | 2 | 8 |
| LL-078 | 2 | 2 | 2 | 0 | 2 | 8 |
| LL-078 | 3 | 1 | 2 | 0 | 2 | 8 |
| LL-078 | 2 | 1 | 1 | 0 | 1 | 5 |
| LL-078 | 2 | 1 | 0 | 0 | 1 | 4 |
| LL-078 | 1 | 1 | 0 | 0 | 1 | 3 |

**Supplementary Table 2.**
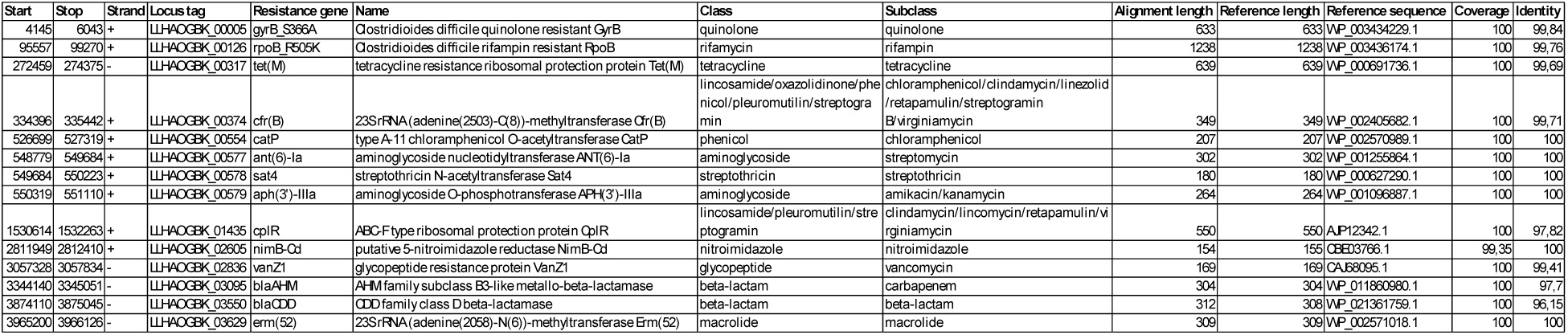
Antimicrobial resistance genes of L-NTCD03.

